# How to select the best model from AlphaFold2 structures?

**DOI:** 10.1101/2022.04.05.487218

**Authors:** Yuma Takei, Takashi Ishida

## Abstract

Among the methods for protein structure prediction, which is important in biological research, AlphaFold2 has demonstrated astonishing accuracy in the 14th Community Wide Experiment on the Critical Assessment of Techniques for Protein Structure Prediction (CASP14). The accuracy is close to the level of experimental structure determination. Furthermore, AlphaFold2 predicts three-dimensional structures and estimates the accuracy of the predicted structures. AlphaFold2 outputs two model accuracy estimation scores, pLDDT, and pTM, enabling the user to judge the reliability of the predicted structures. Original research of AlphaFold2 showed that those scores had good correlations to actual prediction accuracy. However, it was unclear whether we could select a structure close to the native structure when multiple structures are predicted for a single protein. In this study, we generated several hundred structures with different combinations of parameters for 500 proteins and verified the performance of the accuracy estimation scores of AlphaFold2. In addition, we compared those scores with existing accuracy estimation methods. As a result, pLDDT and pTM showed better performance than the existing accuracy estimation methods for AlphaFold2 structures. However, the estimation performance of relative accuracy of the scores was still insufficient, and the improvement would be needed for further utilization of AlphaFold2.

## Introduction

Protein conformational information is important in biological research such as drug discovery[1, 2, 3, 4]. Since the experimental determination of the three-dimensional structure of a protein is time-consuming and costly, the computational prediction has been studied[5, 6]. Recently, various methods have been developed using deep learning, and the prediction accuracies have become higher than before[7, 8, 9, 10, 11]. In the 14th Community Wide Experiment on the Critical Assessment of Techniques for Protein Structure Prediction (CASP14)[12], an international benchmark of protein structure prediction methods, AlphaFold2[13] outperformed all of the other methods. In the CASP14, the median of structure model accuracies in GDT_TS of AlphaFold2 to the native structure was 0.924[14], which could be considered to be comparable to experimental accuracy[15]. Due to its remarkable accuracy, AlphaFold2 has already been used in applied research such as protein-protein interaction (PPI) prediction[16, 17] and structure-based drug design[18], and will continue to be used in the future.

However, there are still several unclear points in the structure prediction using AlphaFold2. One of them is the diversity of predicted structures for the same target. AlphaFold2 has several parameters specified at execution, including the prediction model, number of recycles, random seed, etc. Combining these parameters makes it possible to generate multiple structures for a single protein. A study on PPI prediction using AlphaFold2 revealed that the correctness rate of the complex varies depending on the parameters[16]. Holo -structures are more likely to be output than apo-structures when multiple prediction models are used[19]. These results clearly show that different parameters can generate different predicted structures. However, no studies have yet quantitatively demonstrated the difference in the accuracy of multiple predicted structures generated with different parameters, using a large target set.

Thus, we first generated several hundred structures by combining parameters for each of the 500 protein sequences not included in the AlphaFold2 training and verified the difference in the accuracy of the predicted structures. As a result, for approximately 30% of the targets, AlphaFold2 outputted predicted structures with different GDT_TS[20] more than 0.05. Large prediction accuracy differences (GDT_TS larger than 0.2) were observed for some of the targets. Unfortunately, no perfect parameter setting could output the best predicted structure for any proteins.

Thus, if we want to obtain a better structure model using AlphaFold2, we should generate multiple structures with different parameters and need to select the best one from them to use in the application study. However, in real-world situations, the structure with the highest accuracy cannot be identified because there is no correct answer (the native structure), and we cannot calculate the structural similarity of the structures to the native structures. Additionally, it is unclear whether the accuracy of the predicted structure is sufficient to be used in the subsequent study.

Model quality assessment (MQA) (or estimation of model accuracy, EMA) can be used to estimate the accuracy of structures. MQA generally takes a predicted structure as input and outputs a score that indicates the quality of that structure. To date, statistical potential function-based method[21, 22] and machine learning-based methods[23, 24, 25, 26] have been developed. Recently, deep learning-based methods have been proposed, and the performance became higher than conventional methods[27]. Using a high-performance MQA method, it will be possible to select a structure close to the native structure from multiple structures, and it will be possible to judge the reliability of those structures.

Fortunately, AlphaFold2 not only predicts 3D structures but also estimates the predicted model quality using two different scores, the pLDDT and pTM scores. Those scores allow the user to judge the reliability of the predicted structure. The authors of AlphaFold2 showed that the scores for different proteins have a high correlation with actual prediction accuracy (structural similarity to the native structure)[13, 28]. The results showed those scores could be used to judge the reliability of the predicted structure roughly. Using the score, we can judge whether the predicted structure for a protein A is better than the predicted structure for another protein B. However, it is unclear whether the pLDDT and pTM can be used to select the best structure when multiple structures are generated for a single protein. It is also unclear whether pLDDT and pTM have better model quality estimation performance than the existing MQA methods. Thus, we do not know how to select the best structure model from AlphaFold2 predictions.

In this study, we benchmarked the accuracy estimation performance of pLDDT and pTM from the viewpoint of the best model selection of predicted structures. To test them, we generated a dataset including several hundred structures predicted by AlphaFold2 with different combinations of the parameters for each of the 500 proteins not included in the AlphaFold2 training.

## Materials and methods

In this study, we first generated hundreds of structure models for each of the 500 targets using AlphaFold2 with different combinations of the parameters. Then, we examined how much the prediction accuracy of AlphaFold2 structures varied in each target. Next, we applied existing MQA methods for the predicted structures to evaluate the existing MQA methods and the accuracy estimation scores of AlphaFold2, pLDDT, and pTM.

### Target selection

We selected protein chains whose structures have already been experimentally determined and were not included in the training data of AlphaFold2. The target protein chains were selected based on the test dataset construction method in the AlphaFold2 study[13], with some modifications.

The procedure for target selection was as follows. First, 27,265 PDB entries were retrieved from the Protein data bank (PDB)[29] using the following conditions: initial release date is from 2018-05-01 to 2021-09-30, determined by X-ray crystallography, and the resolution is 2.5 Å or better. The obtained entries were clustered with 40% sequence identity per chain using cluster data published by PDB. We selected 5,937 protein chains from each cluster with the best resolution that satisfied the following conditions: the size is from 80 to 700 residues, not including non-standard amino acids, and the ratio of the missing residue is less than 20%. We divided the dataset into two subsets based on the sequence similarity to AlphaFold2 training data. Of these, 3,217 protein chains were similar to those in the AlphaFold2 training data, and 2,720 protein chains were sequentially dissimilar. To determine whether similar sequences existed in the AlphaFold2 training data, we used the presence of entries released before 2018-04-30 that were used as training data of AlphaFold2 in the clusters with 40% sequence identity. Finally, since it is computationally costly to run AlphaFold2 for all protein chains, we sampled 500 of them and used them as the targets. Of the 500 protein chains, 250 protein chains were randomly sampled from a subset that had similar sequences to the AlphaFold2 training data, and 250 were randomly sampled from a subset that did not have similar sequences to the AlphaFold2 training data. Additionally, we excluded protein structures composed of multiple structural domains, and their interactions were insufficient because they are inappropriate for evaluating the prediction accuracy for the entire structure. The details of the exclusion method are shown in the supplementary method in S1 File.

### AlphaFold2 setting

#### Parameters

By using different combinations of the parameters of AlphaFold2, hundreds of prediction structures are generated per target. In this study, we changed the following parameters: prediction model, number of ensembles, number of recycles, and random seed. As the prediction model, five normal models used in CASP14 and five ptm models fine-tuned to output pTM scores are available. We used all of these ten prediction models. The number of ensembles determines the number of times the multiple sequence alignment (MSA) is sampled. We used one and eight for the number of ensembles. An ensemble number of one means no ensemble. The number of recycles determines the number of times the output structure is iteratively used to input the AlphaFold2’s neural network. For the number of recycles, we used ten recycle numbers from 1 to 10. The random seed sets the seed for the stochastic operations of AlphaFold2. We used two types of random seeds. Finally, by combining these parameters, we generated 400 structures for a single target (= 10 (models) × 2 (ensembles) × 10 (recycles) × 2 (random seed)). However, for targets with more than 400 residues, only ensemble number one was used because ensemble number eight could not be executed due to insufficient GPU memory, and 200 structures were generated for the targets. The optimization by Amber force field[30] was not used because it was almost ineffective in the preliminary verification.

#### Execution environment

Currently, several versions of AlphaFold2 employ different sequence searches and have different functions. In this study, we used LocalColabFold[31] v1.0.0, which enables ColabFold[32] to be used in a local environment. The difference between LocalColabFold and the original one is the MSA generation and template usage. LocalColabFold uses MMseqs2[33, 34] for the MSA generation. In addition, we modified the AlphaFold2 v2.0.1 source code to output all structures during the recycling process. This makes it possible to output the structure of each recycling iteration from 1 to 10 by a single execution (the original implementation only outputs the final structure). The source code is available at https://github.com/yutake27/alphafold.

### Calculation of structure prediction accuracy

To calculate the accuracy of the predicted tertiary structures, we used the GDT_TS[20], GDT_HA, TM-score[35], and lDDT[36]. In this study, we used GDT_TS as the main metric. lDDT evaluates the local structural similarity in residue-level. Thus, we calculate the average of lDDT scores of all residues to evaluate the whole structure. The residues of the model structure and the reference structure must match perfectly when calculating the lDDT. In a few cases, the residues in the amino acid sequence of the protein differ from those in the native structure, and the lDDT cannot be performed on the target. For such targets, the lDDT calculation was omitted.

### Performance evaluation of accuracy estimation

#### Performance evaluation metrics

We evaluated the performance of model accuracy estimation from two different viewpoints. One is the evaluation of the relative accuracy of multiple predicted structures for the same target. This study generated 200 or 400 structures for a single target. This evaluates the ability to select the good models from those structures. The other is the evaluation of the absolute accuracy of a single structure. This evaluates how well the accuracy of a single structure can be estimated.

We used “loss” and “correlation” as measures to evaluate the estimation performance of the relative accuracy of predicted structures for the single target. Loss is the difference between the accuracy of the best structure and the accuracy of the structure selected by an MQA method. The smaller loss is, the better structure is selected among the predicted structures. To judge whether the loss value is good or bad, we compared it with the expected value when the predicted structures are randomly selected. A correlation was used to evaluate the relationship between the actual accuracy of the predicted structure and the accuracy estimated by the estimation method. Pearson correlation and Spearman’s rank correlation were used. All these values were calculated for each target, and finally, the performance for the whole dataset was calculated by averaging.

For evaluating the absolute accuracy of a structure, we used Pearson correlation and mean absolute error (MAE).

#### Benchmarked MQA methods

We evaluated the accuracy estimation performance of MQA methods on AlphaFold2 structures. We checked the performances of pLDDT and pTM, which are outputted by AlphaFold2 prediction to estimate the lDDT and TM-score, respectively. We also compared the performances with those of existing MQA methods. As the existing MQA methods, we used statistical potential function-based methods and machine learning-based methods.

DOPE[21] and SOAP[22] were selected as typical statistical potential function-based methods. DOPE is a potential based on distances between atoms. SOAP is based on the atomic distance and orientation between bonds.

As the machine learning-based methods, we used ProQ3D[37] and SBROD[38], which showed high performance in the accuracy estimation section of CASP13[39], and VoroCNN[40], P3CMQA[41], and DeepAccNet[42], which showed high performance in CASP14[27]. ProQ3D is a multi-layer perceptron-based method that uses sequence profile-based features and rosette energy[43] as input. We used the default S-score version of the ProQ3D prediction models. SBROD is a ridge regression-based method that uses geometric features of the backbone. VoroCNN is a graph convolutional neural network-based method that constructs a graph by Voronoi tessellation of 3D molecular structures. We used the default vorocnn_geometric version. P3CMQA is a three-dimensional convolutional neural network (3DCNN)-based method that uses atomic features and sequence profile-based features as input. DeepAccNet assesses the local atomic environment with a 3DCNN and the global environment with a 2DCNN. There are several versions of DeepAccNet. We used the standard version of DeepAccNet and DeepAccNet-Bert, which adds sequence embedding by ProtTrans[44].

We calculated the scores of all these methods for the regions, excluding the missing residue in the native structure. The average of the scores for the resolved residue was used for methods with local accuracy estimation scores for each residue (ProQ3D, VoroCNN, P3CMQA, DeepAccNet, DeepAccNet-Bert, pLDDT). For DOPE and SOAP, which only have scores for the whole structure, the missing residues in the native structure were excluded from the pdb file of the predicted structure, and then the method was run. In addition, pTM was calculated by excluding missing residues from the Predicted Alignment Error output by AlphaFold2.

## Results and Discussion

### Structure prediction accuracy of AlphaFold2

In this study, AlphaFold2 was run on 500 protein target sequences, and its prediction accuracy was evaluated using the structural similarity labels GDT_TS, GDT_HA, TM-score, and lDDT. The distribution of the maximum value of each label for each target is shown in Fig 1. The median of the maximum GDT_TS for each target was 0.968, and the median of the mean lDDT was 0.899. As already shown in the AlphaFold2 paper, the prediction accuracy of AlphaFold2 is extremely high, and the results are compatible with them[13]. Only 14 of the 500 targets had a GDT_TS of less than 0.8, and 11 had a GDT_TS of approximately 0.7. In the remaining three targets, the GDT_TS was less than 0.5, and the predictions were not successful.

**Fig 1:**
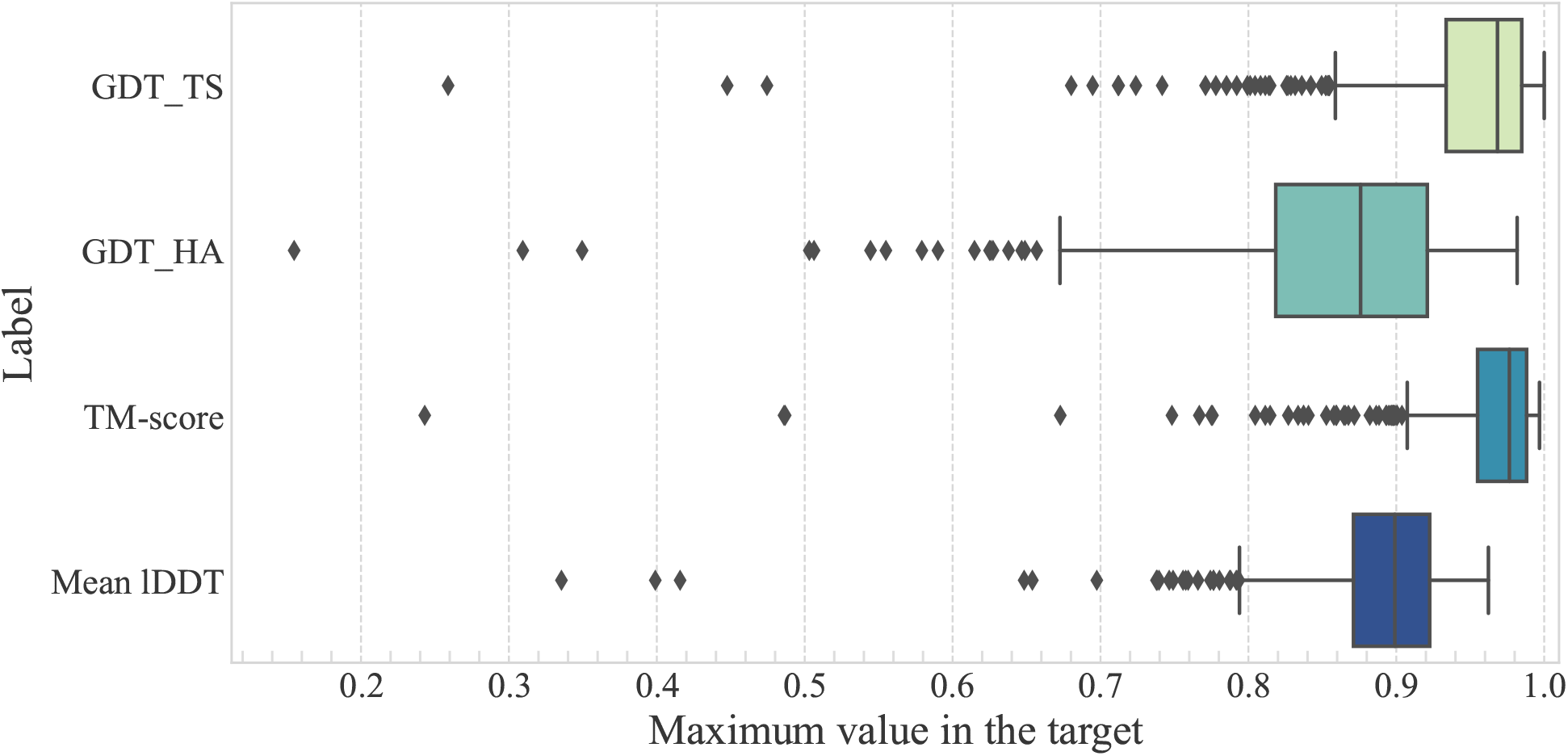
Box plot of the maximum value of labels in each target. Mean lDDT is the average value of lDDT.

We confirmed the native structures of the targets with GDT_TS less than 0.8 and found that 6Z4U_A[45] and 6NEK_A[46] are homo-dimers, and their stable structures are formed by the interaction of two chains. Since it is inappropriate to evaluate the prediction accuracy of such proteins using a single chain, we excluded them from the subsequent evaluation. The native and predicted structures of the targets with low accuracy are shown in Fig S1 in S1 File.

### Difference in the accuracy of predicted structures for the single target

This study generated 200 or 400 structures depending on the sequence length for a single target by combining several parameters. We examined how accuracy differs among these structures for the same target.

The median difference between the maximum and minimum GDT_TS for each target was 0.034. Fig 2A shows the distribution. In about 30% of the targets, the difference was 0.05 or more. In addition, about 10% of the targets had a difference of 0.1 or more, and 21 targets of these had a large difference of 0.2 or more. This result indicates that although not all targets have a difference in accuracy between the predicted structures, some targets do. The distribution of the difference between the maximum and minimum values of each accuracy label and the standard deviation of the labels within a target is shown in Section 2.3 in S1 File.

**Fig 2:**
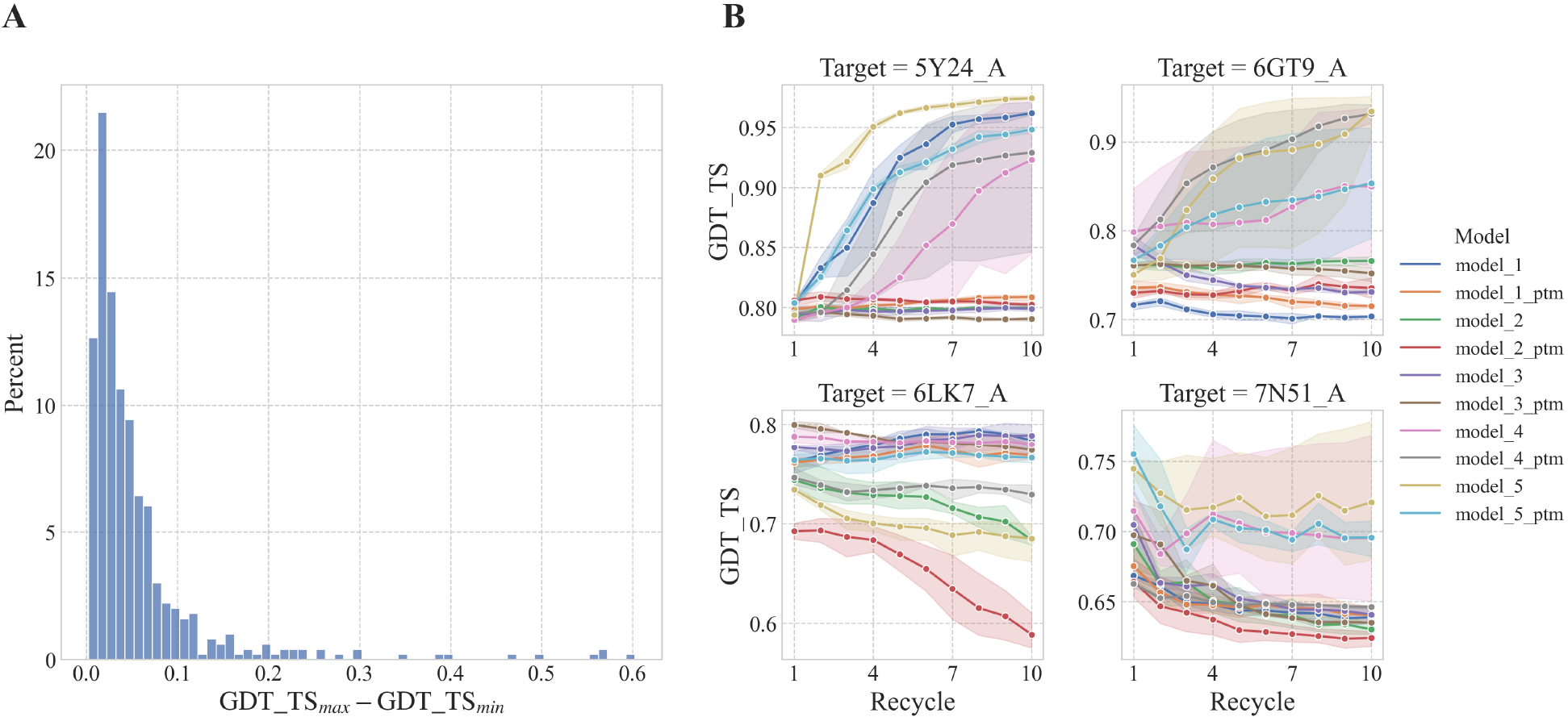
Difference in prediction accuracy within a single target. (A) Histogram of difference between maximum and minimum GDT_TS. (B) Line plot of GDT_TS for each recycling. The color of the line represents the prediction model. The mean value of GDT_TS for the structures with the same number of recycles and the prediction model is represented by a point, and the range represents the 95% confidence interval.

### Variation of accuracy by parameters

This study generated 200 or 400 structures depending on the sequence length for a single target by combining the following parameters: prediction model, number of recycles, number of ensembles, and random seed. We examined the difference in accuracy depending on these parameters.

Table 1 shows the accuracy for specific combination of parameters. The accuracy was higher when the number of recycles was 3 and 10 than when it was one. There was no large difference in accuracy between 3 and 10 recycles. However, for some targets, the accuracy decreased by recycling. Fig 2B shows the examples of cases where recycling was successful and harmful. In addition, the accuracy of the normal model was slightly higher than that of the ptm model. Moreover, there was no significant difference in prediction accuracy between the ensemble and random seeds. The details of the difference in accuracy for each parameter are shown in Section 2.4 in S1 File. Those results indicate that there is no clear best combination of parameters, but, as we showed above, AlphaFold2 outputs structure with different accuracy according to the parameter setting. Thus, selecting a structure from predicted structures is important when we want to obtain a a better structure model using AlphaFold2.

**Table 1:** Average structure prediction accuracy with different combination of parameters.

| Label | GDT_TS |  | GDT_HA |  | TM-score |  | Mean IDDT |  |
| --- | --- | --- | --- | --- | --- | --- | --- | --- |
| Model | normal | ptm | normal | ptm | normal | ptm | normal | ptm |
| 1 recycle | 0.927 | 0.924 | 0.820 | 0.817 | 0.948 | 0.946 | 0.871 | 0.869 |
| 3 recycle | 0.930 | 0.927 | 0.826 | 0.823 | 0.951 | 0.949 | 0.875 | 0.873 |
| 10 recycle | 0.930 | 0.928 | 0.825 | 0.823 | 0.951 | 0.949 | 0.875 | 0.873 |

The first row shows the labels and the second row shows the prediction model. The first column shows the number of recycles. The second and subsequent rows show the mean of the median labels for each target.

### MQA performance comparison on AlphaFold2 structure

We compared the performance of AlphaFold2 confidence scores, pLDDT, and pTM, with that of existing MQA methods. The performance was evaluated in two different viewpoints: the estimation performance of the relative accuracy of structure within the target and the absolute accuracy of the structure. To appropriately compare pTM scores with other MQA methods in the evaluation, we used only the structures predicted from the ptm model with pTM score. Since there was almost no difference in the accuracy between the structures predicted by the normal model and the structures predicted by the ptm model in each target, it is considered acceptable to focus only on the structures predicted by the ptm model.

#### Estimation performance of relative accuracy

Loss, Pearson correlation, and Spearman correlation were used to evaluate the performance of relative accuracy estimation of the predicted structures for each target. We considered that the targets with small differences in the accuracy of the predicted structures within the target are inappropriate for evaluating relative accuracy estimation. Thus, only 150 targets with the difference between the maximum and minimum GDT_TS values greater than 0.05 were used for evaluation.

The average values of loss, Pearson correlation, and Spearman correlation per target for GDT_TS are shown in Table 2. For all metrics, pLDDT or pTM was the best, and there was no significant difference between them. The Wilcoxon signed-rank test against pLDDT was conducted at a significance level of 1% to check if there was a statistically significant difference between pLDDT and the other methods. For loss, pLDDT performed significantly better than random selection. Also, pLDDT was significantly better than some methods for loss, but not for all of the MQA methods. Similarly, Pearson and Spearman correlations were significant for some methods but not for all. For the labels GDT_HA, TM-score, and lDDT, pLDDT or pTM was the best as well (Table S1 in S1 File).

**Table 2:** Estimation performance of relative accuracy.

| Method | Loss | Pearson | Spearman |
| --- | --- | --- | --- |
| DOPE | 3.839 | 0.309 | 0.248 |
| SOAP | 3.248 | *0.279 | 0.236 |
| ProQ3D | *4.712 | *0.174 | *0.123 |
| SBROD | *4.940 | *0.082 | *0.066 |
| VoroCNN | 4.046 | *0.188 | *0.150 |
| P3CMQA | 3.921 | *0.183 | *0.140 |
| DeepAccNet | 3.368 | 0.275 | 0.202 |
| DeepAccNet-Bert | 3.409 | *0.236 | 0.190 |
| pLDDT | 3.084 | **0.360** | **0.279** |
| pTM | **2.939** | 0.353 | 0.270 |
| Random selection | *4.556 | - | - |

The first column shows the method name. The second to fourth columns represent the average value of loss, Pearson correlation, and Spearman correlation for each target, respectively, when GDT\_TS is used as a label. The loss values are multiplied by 100 for clarity. An asterisk means that the p-value was less than 0.01 when conducting the Wilcoxon signed-rank test against pLDDT.

Although pLDDT and pTM were the best for all metrics, their estimation performance was insufficient, and they failed significantly for some targets. In terms of loss, pLDDT and pTM had about 10 targets whose loss was farther than 10. For the Pearson and Spearman correlations, about 25% of the targets were negatively correlated, and the average values of pLDDT were as low as 0.360 and 0.279, respectively. Thus, the relative accuracy of the structures in each target was not well recognized. The distribution of each metric per target is shown in Fig S11 in S1 File.

Among the existing MQA methods, DeepAccNet, DOPE, and SOAP showed high performance. Although DOPE and SOAP are classical methods based on statistical potential functions derived from native structures, their performances were higher than that of machine learning-based methods. This may suggest that learning the distribution of native structures effectively estimates the accuracy of highly accurate 3D structures. Generally, machine learning-based MQA methods showed better performance than potential-based methods in CASP[27], but the results showed the opposite. The problem of training data may be one of the reasons. Most machine learning-based methods use the CASP dataset[12] as training data, and the number of native structures used for training is only about 300. DeepAccNet is a machine learning-based method, but it uses about 7,000 native structures for the training. There is a large difference in the number of native structures between DeepAccNet and CASP datasets. In addition, the CASP dataset does not contain many highly accurate structures such as those predicted by AlphaFold2. This lack of native structures and highly accurate structures in the training data would be a reason for the poor performance of machine learning-based methods except for DeepAccNet.

#### Estimation performance of absolute accuracy

We used Pearson correlation and MAE to evaluate the estimation performance of absolute accuracy. Only methods with scores normalized from 0 to 1 were used for evaluation.

A scatter plot between GDT_TS and the scores of each MQA method is shown in Fig 3. pLDDT was the best in both Pearson correlation and MAE, and was better than pTM for GDT_TS. Among the other MQA methods, DeepAccNet was the best, but its correlation was much lower than that of pLDDT.

**Fig 3:**
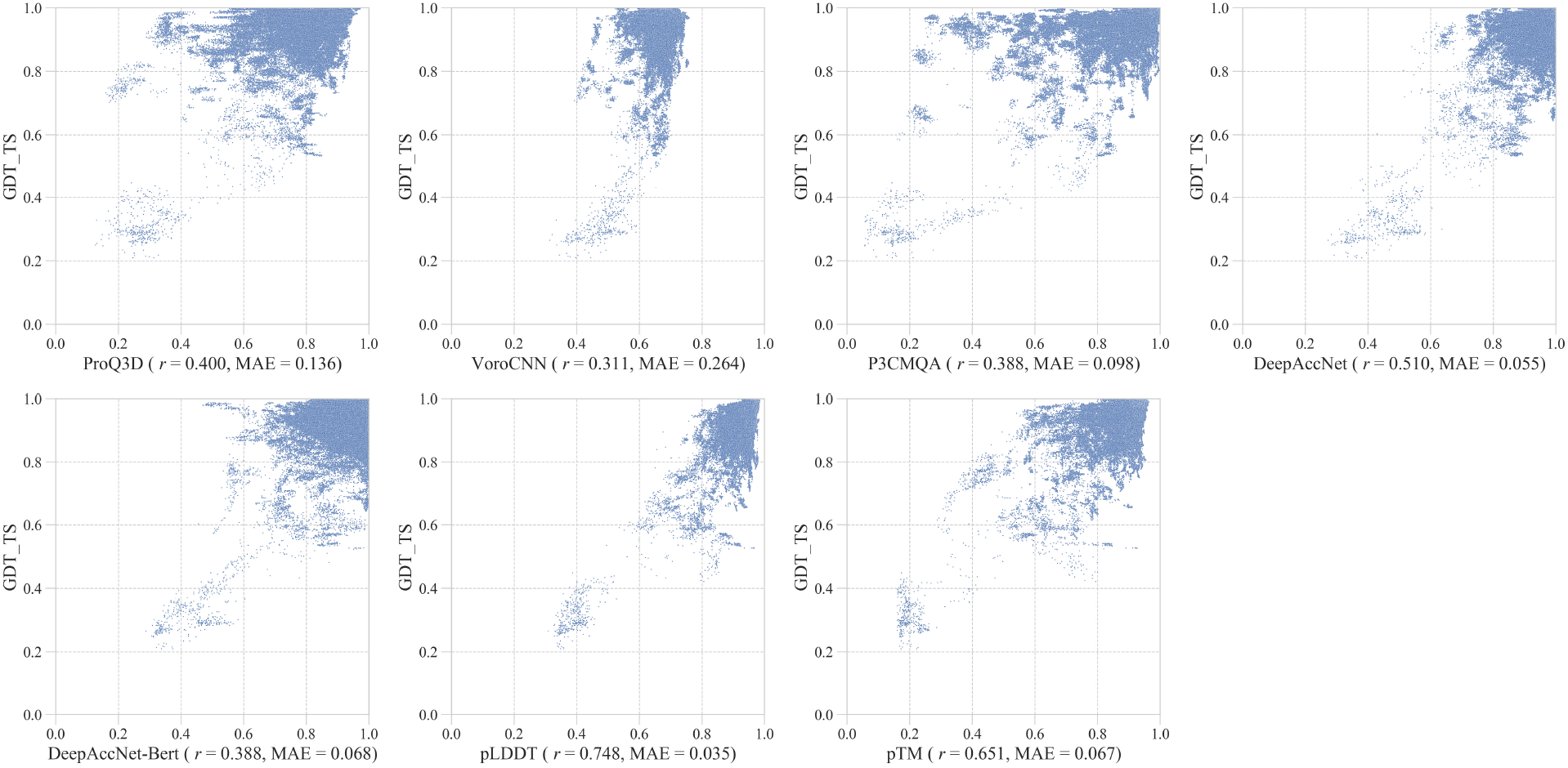
Scatter plot between GDT TS and MQA methods.

The results for the mean lDDT and TM-score are shown in Fig S12 and S13 in S1 File, respectively. Similar to the results for GDT_TS, the correlations of pLDDT and pTM are much higher than those of other MQA methods.

### Targets with low structure prediction accuracy

As shown in the results, most of the predicted structures of AlphaFold2 had sufficient accuracy. However, some targets could not be predicted with high accuracy. The original AlphaFold2 research indicated the prediction accuracy depends on the number of similar sequences of the target[13]. Thus, we also tried to analyze the factors related to the low accuracy.

We first checked the relationship between the number of effective sequences (Neff) in the MSA and the accuracy. Neff was computed following the previous study[47, 16]. Scatter plots between Neff and GDT_TS are shown in Fig 4A. A larger Neff increases the percentage of targets with higher GDT_TS. However, there are more targets with GDT_TS greater than 0.8 than those with GDT_TS less than 0.8 even when Neff is less than 10. Thus, although Neff is a good indicator to estimate the accuracy of a prediction of AlphaFold2, the structure prediction accuracy cannot be determined only by Neff.

**Fig 4:**
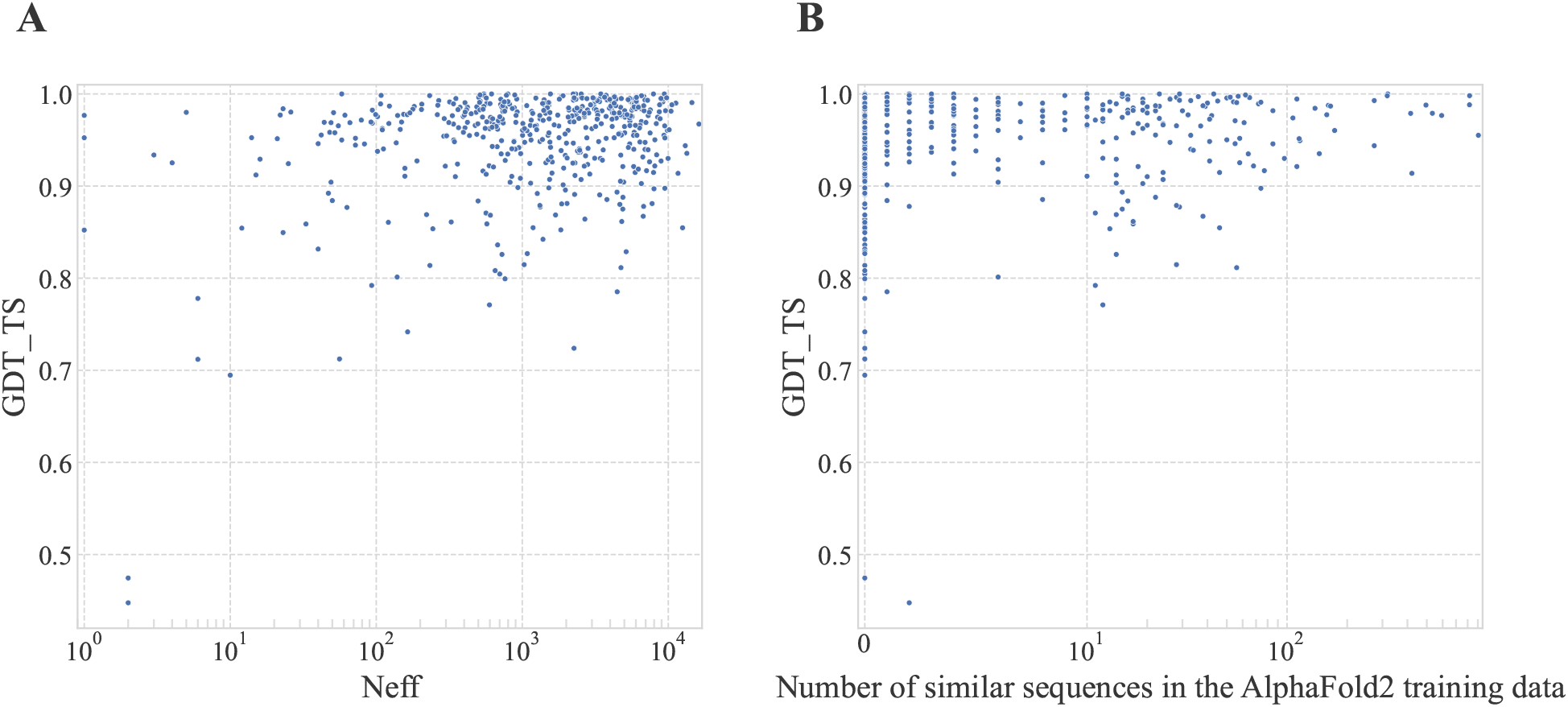
Relationship between structure prediction accuracy and the properties of the target. (A) Scatter plot between Neff and GDT_TS. (B) Scatter plot between number of similar sequences in AlphaFold2 training data and GDT_TS. The number of similar sequences was derived from the cluster size of 40% sequence identity clustering for PDB entries released up to the maximum release date of the AlphaFold2 training data.

Next, we analyzed the relationship between the number of similar sequences in the AlphaFold2 training set and the accuracy. Fig 4B shows a scatter plot between the number of similar sequences in the AlphaFold2 training data and GDT_TS. As the number of similar sequences decreases, the number of targets with low accuracy increases. The mean values of the maximum GDT_TS for targets with and without similar sequences are 0.956 and 0.943, respectively. The Mann-Whitney U test was used to test the difference in GDT_TS at a significance level of 1%, and it was significant (P-value = 7.44 × 10^−4^). Thus, there is a tendency for the accuracy to decrease when similar sequences are not included. However, even when there were few or no similar sequences, some targets were successfully predicted with a GDT_TS of 0.9 or higher. The number of similar sequences is considered an important factor related to the prediction accuracy, but there are cases where the prediction is successful even when there are few similar sequences.

Unfortunately, we could not find any other factors that relate to the prediction accuracy. The elucidation of the other factors to estimate the prediction accuracy is still important to use AlphaFold2 better.

### Analysis of the factors causing the accuracy differences within a target

When generating structures for the same target with different parameters, there were sometimes significant differences in the accuracy for some targets. We then analyzed the targets that had the prediction accuracy differences because of the different parameters of the prediction.

First, we checked the difference in the variation of prediction accuracy within a target between two subsets, with similar sequences and dissimilar sequences in the AlphaFold2 training data. The median difference between the maximum GDT_TS and minimum GDT_TS for targets with similar sequences was 0.025, whereas it was 0.043 for targets with dissimilar sequences. The difference between the two was tested using the Mann-Whitney U test with a significance level of 1% and was statistically significant (P-value = 6.71 × 10^−10^). Thus, we should optimize prediction parameters more when the prediction targets do not have similar training data sequences. The distribution of the difference in accuracy with and without similar sequences in the training data for each label is shown Fig S14 in S1 File.

Next, we tested the difference in the variation of prediction accuracy by the domain number of the target protein. The domain number of each target was obtained from the ECOD database[48, 49] classification (version 2021-10-04). The median difference between the maximum and minimum GDT_TS for single-domain targets was 0.029, compared to 0.044 for multi-domain targets. The difference between the two was tested using the Mann-Whitney U test with the significance level at 1% and was significant (P-value = 2.13 × 10^−5^). The multi-domain target is more likely to have a difference in accuracy than the single-domain target. The distribution of the difference in prediction accuracy by domain number is shown Fig S15 in S1 File.

### Detection of low accuracy predictions

In the results section, we evaluated the performance of the MQA method based on the performance that the method can find the best structure from predicted structures. However, AlphaFold2 sometimes failed to predict accurate structures, and thus, it is also important to detect cases that AlphaFold2 fails to predict. To check this point, we tested the performance of MQA methods using the following two metrics. We classified the predicted structures into successed ones (positive) and failed ones (negative) based on the prediction accuracy and a threshold. The first metric is the Area Under the ROC Curve (AUC) calculated from predicted structures of multiple targets. It evaluates the detection performance of absolutely low accuracy structures, regardless of the target. The second is the AUC, calculated from predicted structures of a single target. This evaluates the performance of classifying relatively accurate and inaccurate structures within a target.

First, we verified the detection performance of absolutely low-accurate structures among the predicted structures. To define negative examples based on the absolute accuracy of the predicted structure, we used the definition of a threshold for outliers in a general box plot (first quartile minus 1.5 times the interquartile range) with the label GDT_TS. Using this definition, the threshold for negative examples was GDT_TS 0.807. Based on this threshold, positive and negative examples were labeled, and the AUC was calculated. The results are shown in Fig 5A. The AUC for pTM was the best, and pLDDT was comparable to pTM. Other MQA methods showed better performances than random selection, but they were inferior to pLDDT and pTM. Similar results were obtained when we used the different thresholds, 0.6 and 0.7. Thus, pLDDT and pTM were also good for detecting structures with absolutely low accuracy.

**Fig 5:**
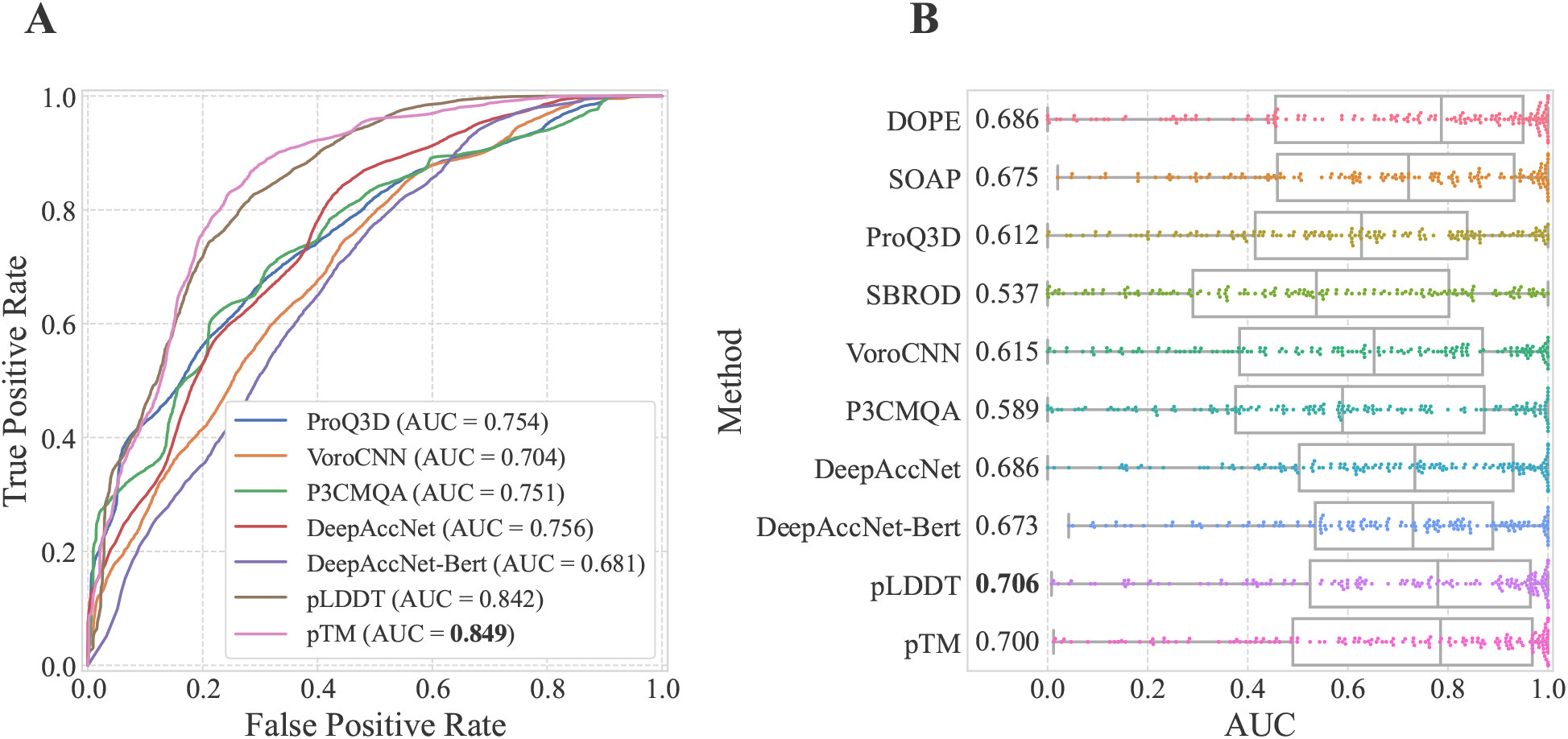
Detection performance of low accurate structures. (A) ROC curve when positive and negative were defined by absolute GDT_TS value. Only structures generated from ptm models were used. In total, there were 87, 933 positive samples and 5, 667 negative samples. Only methods with normalized scores were used for comparison. (B) Distribution of AUC for each target when positive and negative were defined based on relative GDT_TS. Only structures generated from ptm models were used. Only targets with a difference of 0.05 or more in GDT_TS were used.

Next, we examined the detection performance of predicted structures with relatively low accuracy within a target. For each target, Maximum GDT_TS — 0.05 was set as the threshold value for the positive and negative cases, and the AUC was calculated. Fig 5B shows the distribution of AUC for each MQA method. pLDDT was the best, and pTM was comparable to pLDDT. However, there were no significant differences between pLDDT, DOPE, and DeepAccNet. Even using pLDDT, the AUC was approximately 0.7, and 25% of targets showed the AUC less than 0.5. Thus, identifying relatively accurate predictions within a target was still difficult, and the performance should be improved.

### Analysis of the factors that cause AlphaFold2 to fail in accuracy estimation

The MQA performances of the accuracy estimation scores of AlphaFold2, pLDDT, and pTM, were better than those of the other existing MQA methods, but the accuracy estimation failed substantially for some targets. Thus, we analyzed to find the factors that cause failures of the accuracy estimation.

We checked the Neff of MSA, and the number of similar sequences in the AlphaFold2 training data was the same as the analysis of AlphaFold2 prediction accuracy. The smaller the Neff, the less available evolutionary information and accuracy estimation may fail. If there are few similar sequences in the training data, the accuracy estimation may fail because similar sequences have not been learned. Thus, we checked the correlation between the MQA performance of pLDDT and Neff (Fig S16 in S1 File) and the correlation between the number of similar sequences and the MQA performance of pLDDT (Fig S17 in S1 File). Unfortunately, there is no correlation in both cases.

Next, we examined the targets for which pLDDT and pTM failed to select the best structure (GDT_TS loss > 0.1). The number of targets with a GDT_TS loss greater than 0.1 for both pLDDT and pTM was 10. Seven of the targets were common. This suggests that pLDDT and pTM fail selection on similar targets. Furthermore, we checked the three-dimensional structures of the best structure and the selected structure by pLDDT and pTM in these targets. We found that there were two types of selected structures (Fig S21, S22, S23, and S25 in S1 File): structures that succeeds in rough prediction, yet the details were off and the prediction accuracy was low and structures that have high prediction accuracy for individual domains, yet fails to predict the relative positions between domains. Since pLDDT is originally a score that estimates the prediction accuracy around each residue, it is difficult to estimate the accuracy of the global structure, such as the positional relationship between domains. pTM predicts the accuracy of the entire structure and is considered more likely to detect relative positional shifts between domains than pLDDT, but it failed to select a common target with pLDDT.

Targets with multiple domains, which pLDDT and pTM failed to select, were not excluded during target selection because there is a certain amount of interaction between domains. However, some targets have interdomain flexibility; thus, the structures selected by pLDDT and pTM were not necessarily the structures with low accuracy in structure prediction. Thus, we evaluated estimation performance of relative accuracy only for single-domain targets. We used the classification of domains from the ECOD database. There were 333 single-domain targets out of 500, of which 77 targets with a GDT_TS difference of 0.05 or more were used in the evaluation. The distribution of loss for each MQA method is shown in Fig 6. The loss of pLDDT and pTM was about 0.02, and there were only two targets that failed in selection significantly. Therefore, the performance of pLDDT and pTM in selecting the best structure for single-domain targets is sufficiently high, while the selection of the best structure for multi-domain targets often fails. The detailed results for the analysis are shown in Table S2 in S1 File.

**Fig 6:**
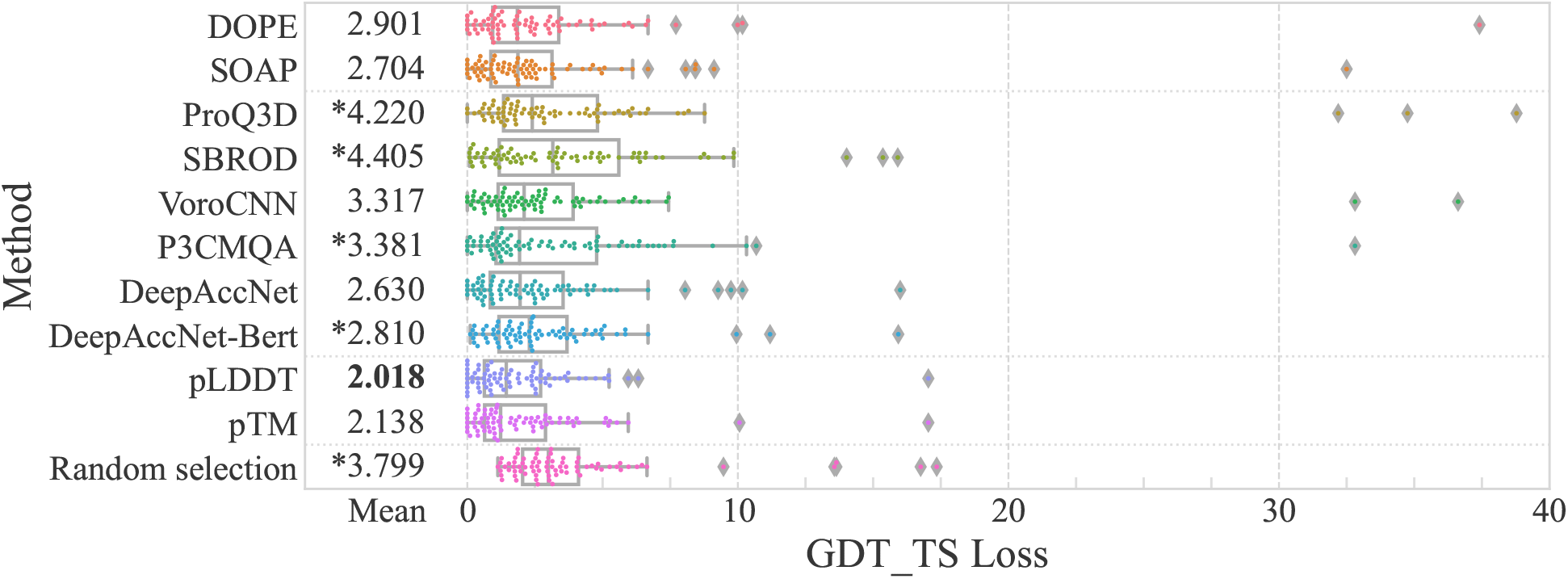
Distribution of GDTTS loss for the single-domain target. The loss values are multiplied by 100 for clarity.

## Conclusion

This study evaluated the accuracy estimation performances of MQA methods for AlphaFold2 structures. We constructed a dataset containing 500 proteins not included in the AlphaFold2 training data and generated several hundred structures using AlphaFold2 with different combinations of parameters. The structure prediction accuracy of AlphaFold2 for the dataset was high, and the median maximum GDT_TS was 0.968. However, there were some targets with lower accuracy (GDT_TS < 0.8), accounting for about 3% of the targets. Approximately 30% of the targets showed differences in the accuracy of the predicted structures (GDT_TS_*max*_ — GDT_TS_*min*_ > 0.05) by different combination of parameters. No best parameter setting can be used for any targets, indicating the necessity of predicted structure selection.

We then evaluated the accuracy estimation performance of existing MQA methods and the accuracy estimation scores pLDDT and pTM that AlphaFold2 outputs during 3D structure prediction. We evaluated the performance from two perspectives: estimation performance of relative accuracy within a target and estimation performance of absolute accuracy. As a result, pLDDT or pTM performed better than existing MQA methods in both. However, even using pLDDT and pTM, the performances of the relative accuracy estimation for a single target were insufficient. This suggests that further improvement of MQA for AlphaFold2 is required. Furthermore, pLDDT and pTM are outputted when AlphaFold2 predicts 3D structures. Thus, they cannot be used to estimate the accuracy of structures predicted by the other structure prediction methods. It is impossible to compare the accuracy of predicted structures using pLDDT and pTM if a structure prediction method with high accuracy equivalent to AlphaFold2 appears. Therefore, it will be necessary in the future to develop accuracy estimation methods that can select structures with relatively high accuracy from those generated by highly accurate structure prediction methods such as AlphaFold2.

## Data availability

Structural data predicted by AlphaFold2 and the source code are available at https://github.com/yutake27/MQAB4AF2. Entries released from 2018-05-01 to 2021-09-30 in the PDB were used as targets. The PDB SearchAPI and DataAPI were used to fetch target candidates. 40% sequence clustering data was obtained from https://cdn.rcsb.org/resources/sequence/clusters/bc-40.out. The ECOD database (2021-10-04) was used to obtain domain information for the target.

## Supporting information

**S1 File. Supporting information** All supporting information is included in this file.

## Acknowledgments

Numerical calculations were carried out on the TSUBAME3.0 supercomputer at Tokyo Institute of Technology. Part of this work is conducted as research activities of AIST - Tokyo Tech Real World Big-Data Computation Open Innovation Laboratory (RWBC-OIL). The authors would like to thank MMseqs2 team for allowing us to use their web server to generate multiple sequence alignments.

